# Sex-limited diversification of the eye in *Heliconius* butterflies

**DOI:** 10.1101/2022.04.25.489414

**Authors:** Nathan P. Buerkle, Nicholas W. VanKuren, Erica L. Westerman, Marcus R. Kronforst, Stephanie E. Palmer

## Abstract

Butterflies have evolved an immense diversity in eye organization to support a range of vision-based behaviors including courtship, oviposition, and foraging. This diversity has been surveyed extensively across the butterfly phylogeny, and here we take a complementary approach to characterize the eye within a group of closely related *Heliconius* butterflies. Using a combination of immunostaining for different opsins and eyeshine for determining the distribution of light-filtering screening pigments, we identified several sexually dimorphic features of eye organization where male eyes varied and female eyes did not. Ultraviolet (UV) sensitive photoreceptors varied in which of two UV opsins were expressed, including co-expression of both within single photoreceptors, and these differences were consistent with a role in courtship and conspecific identification. Additional differences across species and sex included the distribution of three ommatidial types defined by the expression pattern of UV and blue opsins, the distribution of a red screening pigment, and which ommatidial types expressed the red screening pigment. We hypothesize that female eyes are optimized for a dimorphic behavior such as oviposition, while male eyes adapt to other selective pressures such as the local light environment.

## Introduction

The organization of peripheral sensory systems plays an important role in behavior by specifying what environmental information is available to an animal (Wehner, 1987). Compared to downstream neural circuits, these peripheral locations are an evolutionarily labile target for adaptation, allowing for potentially rapid changes that support behavioral evolution (Bendesky and Bargmann, 2011). The selective pressures that promote peripheral evolution are diverse, including factors that influence foraging, courtship, and oviposition. For example, the evolution of trichromatic color vision in primates likely functions to improve the detection of ripe fruit (Melin et al., 2017, 2013; Regan et al., 2001), while the *Drosophila sechellia* olfactory system is specialized for the detection of its noni plant host (Auer et al., 2020). For courtship, the evolution of the cichlid visual system (Seehausen et al., 2008; Terai et al., 2006) and *Heliothis* moth olfactory system can drive reproductive isolation and speciation (Gould et al., 2010; Lee et al., 2016), while sexually dimorphic plumage in warblers appears to co-evolve with sexually dimorphic visual systems (Bloch, 2015). Lastly, rather than specific behavioral contexts, peripheral systems can evolve to match the statistics of the natural environment (Lythgoe, 1979). These differences are commonly observed in the visual systems of aquatic animals living at different water depths (Fasick and Robinson, 2000; Fuller et al., 2003; Torres-Dowdall et al., 2017) or in birds that prefer different forest strata. Thus, the periphery can respond to a multitude of selective pressures to support a range of adaptive behaviors.

The visual system of butterflies presents an interesting system to explore how different selective pressures affect the organization of the eye. Butterflies have exceptional color vision that plays a prominent role in many of their behaviors. The ancestral eye likely comprised ultraviolet (UV), blue (B), and long wavelength (LW) sensitive opsins organized into ommatidia that each house nine photoreceptors (R1-R9, Fig. 1)(Briscoe, 2008). The R3-8 photoreceptors express the LW opsin, the small R9 cell has a generally unknown opsin expression, and the combination of UV and blue opsins in R1 and R2 defines three ommatidial types (UV-UV, B-B, and UV-B) that tile the eye. This photoreceptor composition is sufficient to support trichromatic color vision, and the common evolution of a red sensitive photoreceptor (Blackiston et al., 2011; Briscoe and Chittka, 2001; Frentiu et al., 2007; Zaccardi et al., 2006) can further support tetrachromatic vision (Koshitaka et al., 2008; Vorobyev and Osorio, 1998). Interestingly, however, this speciose group of insects has evolved an immense diversity in eye organization, with different species having up to fifteen unique photoreceptor types with different spectral sensitivities (Arikawa, 2003; Chen et al., 2016, 2013). This diversity encompasses duplications of all three opsins (Arikawa et al., 2005; Briscoe et al., 2010; Frentiu et al., 2007) as well as the use of screening pigments (Arikawa and Stavenga, 1997; Stavenga, 2002) that function as intraocular filters that absorb and prevent some wavelengths of light from reaching the photoreceptors.

**Figure 1.**
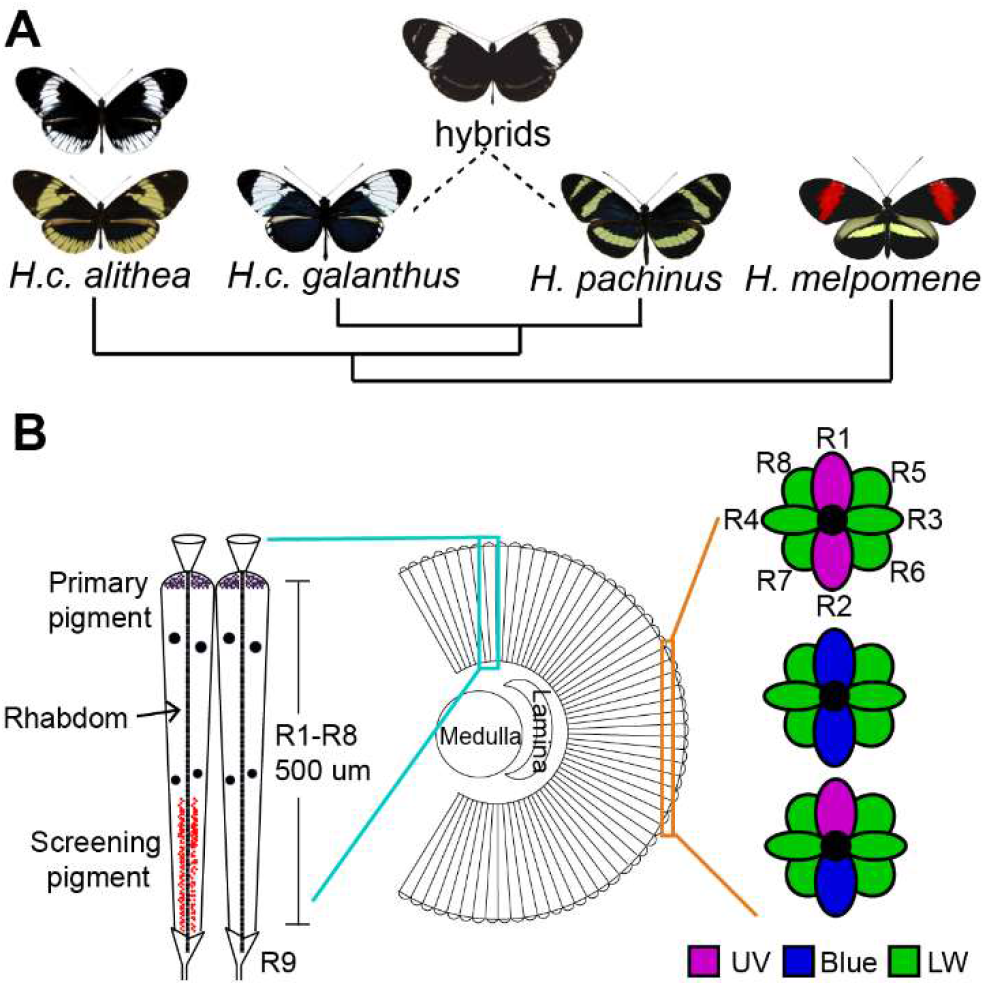
Study system. (A) Phylogenetic tree shows butterflies examined in this study, including the hybrid offspring of two sister species. Wing color is known to be a Mendelian trait, with H.c. alithea being polymorphic. (B) Diagram of basic eye organization shows the anatomy of individual ommatidia. The left shows a longitudinal view of two ommatidia. Note the screening pigments in the proximal portion of the ommatidia, which selectively absorbing certain wavelengths of light to shape photoreceptor spectral sensitivity. The right shows three ommatidia in cross-section, with three ommatidial types defined by which opsins are expressed in the R1 and R2 photoreceptors.

This diversity in eye organization across the butterfly phylogeny highlights the evolvability of the periphery, but the extent to which the eye can evolve among closely related taxa remains unclear. Different aspects of eye organization may vary in their evolvability, such as the number of ommatidial types, photoreceptor spectral sensitivities, or the ratios of different photoreceptor types. Contrasting phylogeny-wide data with comparisons of closely related groups would help distinguish which aspects are especially evolutionarily labile. Thus, in this study we have focused on characterizing eye organization in a group of closely related *Heliconius cydno* butterflies. This genus of mimetic, Neotropical butterflies have a duplicated UV opsin (UV1 ~355 nm, UV2 ~390 nm), a blue opsin (~450 nm), a LW opsin (~550 nm), and a red opsin derived from the LW opsin and a red screening pigment (Briscoe et al., 2010; McCulloch et al., 2016; Zaccardi et al., 2006). A recent study sample 14 species from each of the major *Heliconius* clades (McCulloch et al., 2017), finding substantial differences in eye organization. This survey detected six distinct retinal mosaics defined by the different ommatidial types tiling the eye (e.g. UV1-UV1, UV2-B), ranging from three to six ommatidial types per species as well as several sexual dimorphisms.

In the *Heliconius cydno* butterflies we examined here (Fig. 1), males preferentially court females based predominantly on visual perception of a wing color that has Mendelian inheritance (Westerman et al., 2018), with white wings dominant to yellow. *H.c. galanthus* and *H. pachinus* are white and yellow sister species, respectively, and both strongly prefer to mate with females with the same wing color, while their hybrid offspring court both colors equally (Kronforst et al., 2006). In the polymorphic *H.c. alithea*, yellow males prefer yellow females, while white males court both colors equally (Chamberlain et al., 2009). Thus, understanding how the eye is organized in these taxa is an important first step towards understanding the mechanisms that underlie this divergent behavior that has a simple genetic basis (Van Kuren et al., 2022). We compared eye organization across these taxa as well as the closely related *Heliconius melpomene rosina* using a combination of opsin immunostaining and eyeshine to assay screening pigments. We observed the same basic set of three ommatidial types across all butterflies suggesting the overarching organization of the eye may be phylogenetically constrained. However, we also detected significant variability in the organization of male eyes, but not females, suggesting sex-limited diversification of the eye may be one way that the periphery is able to respond to different selective pressures.

## Results

To characterize the organization of the eye, we first asked which ommatidial types each butterfly expressed by performing antibody staining against UV1, UV2, and blue opsins in thin cross sections of the eye (Fig. 2). Consistent with a previous report (McCulloch et al., 2017), every butterfly had a combination of UV-UV, B-B, and UV-B ommatidia (Fig. 2A). However, the specific UV opsin that was expressed differed with species and sex (Fig. 2A). First, females always expressed only UV1 regardless of taxon, but male eyes varied. *H.c. galanthus* males also expressed UV1, while its sister *H. pachinus* instead expressed UV2. Interestingly, the hybrid offspring of this pair had an intermediate phenotype, showing co-expression of both UV1 and UV2 within single photoreceptors. *H.c. alithea* males of both wing colors always expressed UV2. For *H. melpomene*, we also observed co-expression of both UV1 and UV2 in all males. Surprisingly, qPCR showed within group variability in the degree of co-expression across all groups (Fig. 2B), and we similarly observed co-expression of UV1 and UV2 in three of nine *H.c. galanthus* and four of fifteen *H.c. alithea* (Fig. 2C).

**Figure 2.**
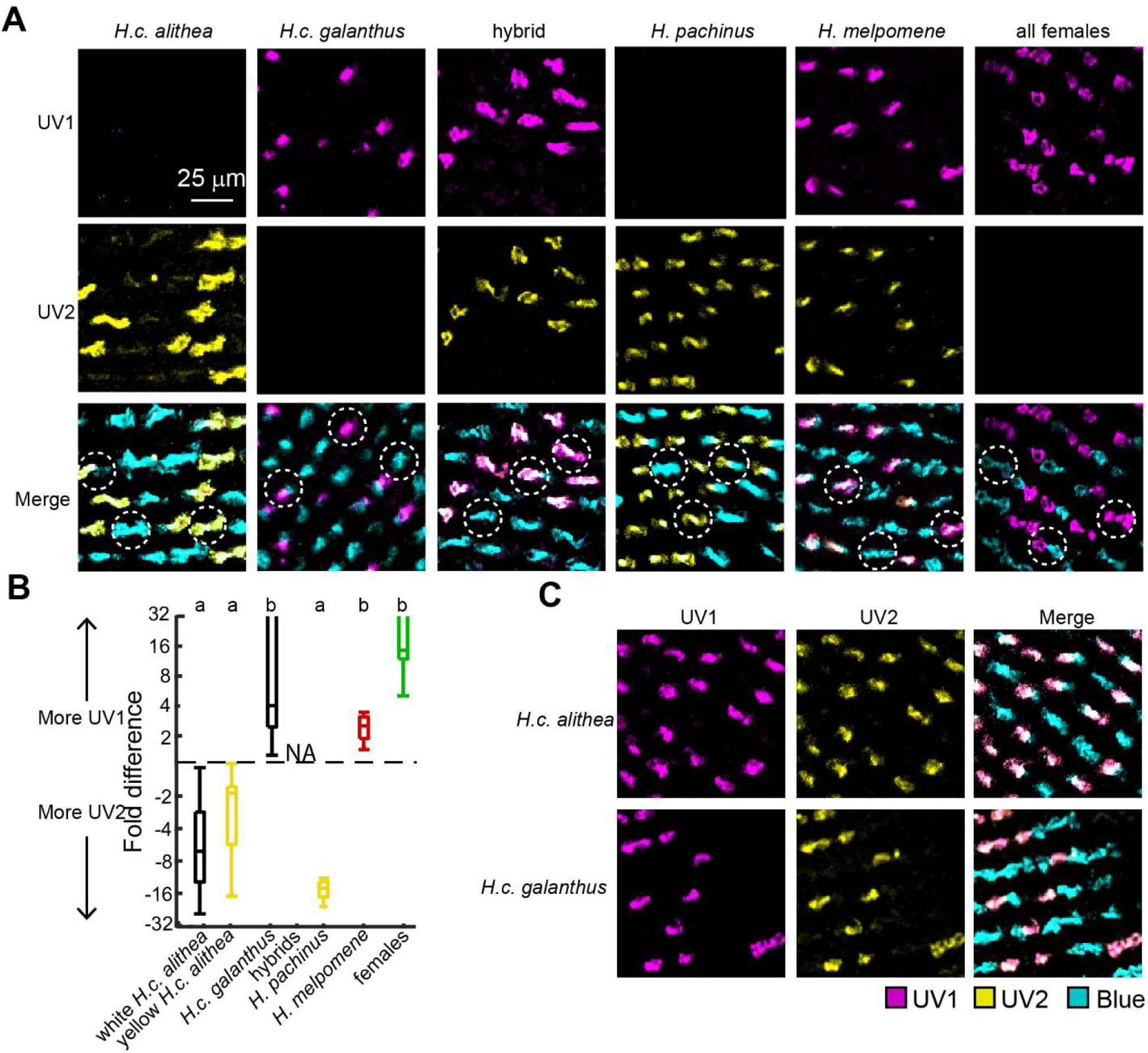
Variable co-expression of two different UV opsins. (A) Antibody staining for UV1, UV2, and blue opsins in thin cross sections of the eye. The first five columns are representative of males in each group, while the sixth column is representative for all females regardless of taxon. This particular female was an *H.c. alithea*. In the bottom row, examples of the three ommatidial types (UV-UV, B-B, and UV-B) are circled. (B) qPCR shows the relative expression levels of UV1 and UV2 across groups on a log scale. The two bars extending off the axis are due to individuals with no detectable UV2 expression. Each bar includes 12 individuals, but no hybrids were used. Letters above indicate groups that are significantly different from each other. (C) Approximately one third of *H.c. alithea* and *H.c. galanthus* males exhibited co-expression of UV1 and UV2. Shown are two representative examples.

We were next interested in determining the distribution of these different ommatidial types across the eye. Since no butterfly had separate UV1 and UV2 photoreceptors, we collectively referred to the expression of any UV opsin as a UV photoreceptor, and therefore counted the number of UV-UV, B-B, and UV-B ommatidia in each immunostained section. We first observed a clear dorsal-ventral difference where the dorsal ~25% of the eye was mostly UV-UV in all butterflies, and the ventral eye had a more even mix of all three ommatidial types (Fig. 3A). In these ventral slices, we counted an average of 500.1 ± 222.0 ommatidia per individual. Figure 3B shows the resulting distributions, and we used hierarchical clustering to assess whether there were any differences across species and sex (Fig. 3C).

**Figure 3.**
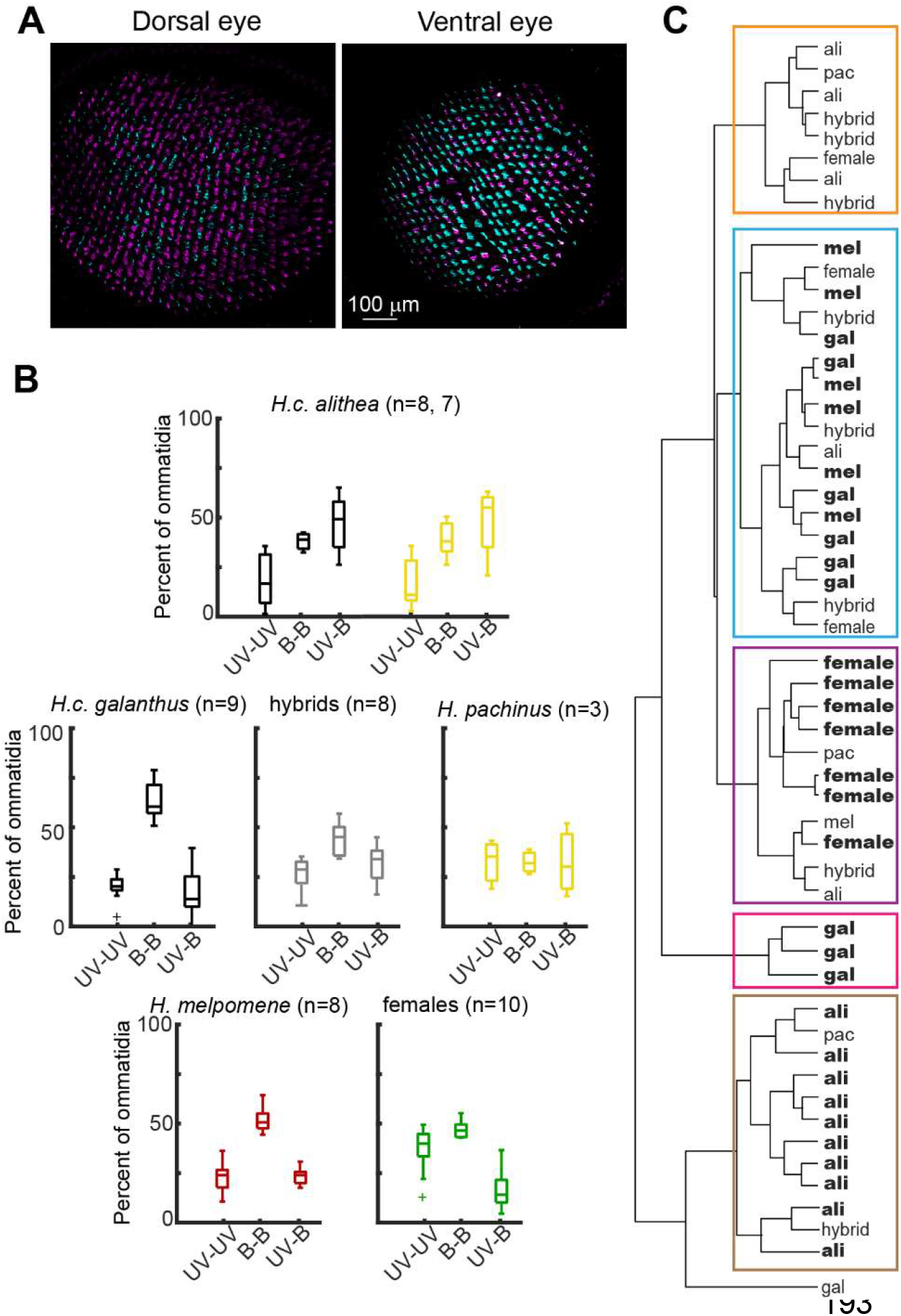
Distributions of the three ommatidial types. (A) Example antibody staining shows differences in the relative proportion of UV and blue photoreceptors in the dorsal and ventral part of the eye. (B) Distributions of the three ommatidial types in the ventral eye. Panels labeled with taxon names are for males only, while all females are grouped together in a single panel. *H.c. alithea* males were separated into two groups based on wing color. (C) Hierarchical clustering of all individuals depicted in panel b. Each of the five major clusters is highlighted with a box, and bolded names show which taxa comprise the majority of each cluster.

Hierarchical clustering detected five major clusters, four of which were predominately populated by one or two groups (Fig. 3C). The most distinct cluster primarily included *H.c. alithea* males of both wing color, which is consistent with visual inspection showing it was the only taxon with mostly UV-B ommatidia (Fig. 3B). *H.c. galanthus* males were split between two clusters, both characterized by eyes with mostly B-B ommatidia. One of these clusters also included most of the *H. melpomene* males, while the second was three males with especially low numbers of UV photoreceptors. Likely due to small sample size, its sister species *H. pachinus* did not cluster together but appeared to have relatively equal numbers of all three ommatidial types. Hybrids also did not cluster together, but similar to UV opsin expression, the distribution appeared to be an average of the distributions observed for the parent species. Finally, we again observed a sexual dimorphism where females of all taxa clustered together and had eyes with few UV-B ommatidia and equal amounts of UV-UV and B-B ommatidia.

In contrast to the diverse expression of UV and blue opsins in the R1 and R2 photoreceptors, the R3-8 photoreceptors all express the green sensitive LW opsin. However, *Heliconius* butterflies also have red sensitive photoreceptors derived from a combination of the LW opsin and a red screening pigment (McCulloch et al., 2016). To assess the distribution of green and red sensitive photoreceptors, we performed eyeshine experiments that reveal which ommatidia express this red pigment. We imaged the eyeshine across the entire dorsal-ventral axis of the eye (Fig. 4), averaging 3,686.9 ± 758.2 ommatidia per butterfly, which is approximately 30% of a *Heliconius* eye (Seymoure et al., 2015). Images from the ventral part of the eye had nearly twice as many ommatidia per photo compared to the rest of the eye (476.8 ± 124.8 vs. 242.5 ± 37.7, p < 0.001). This difference reflects an increase in the spatial resolution of the ventral eye compared to the middle and dorsal part of the eye (Stavenga et al., 2001; Takeuchi et al., 2006).

**Figure 4.**
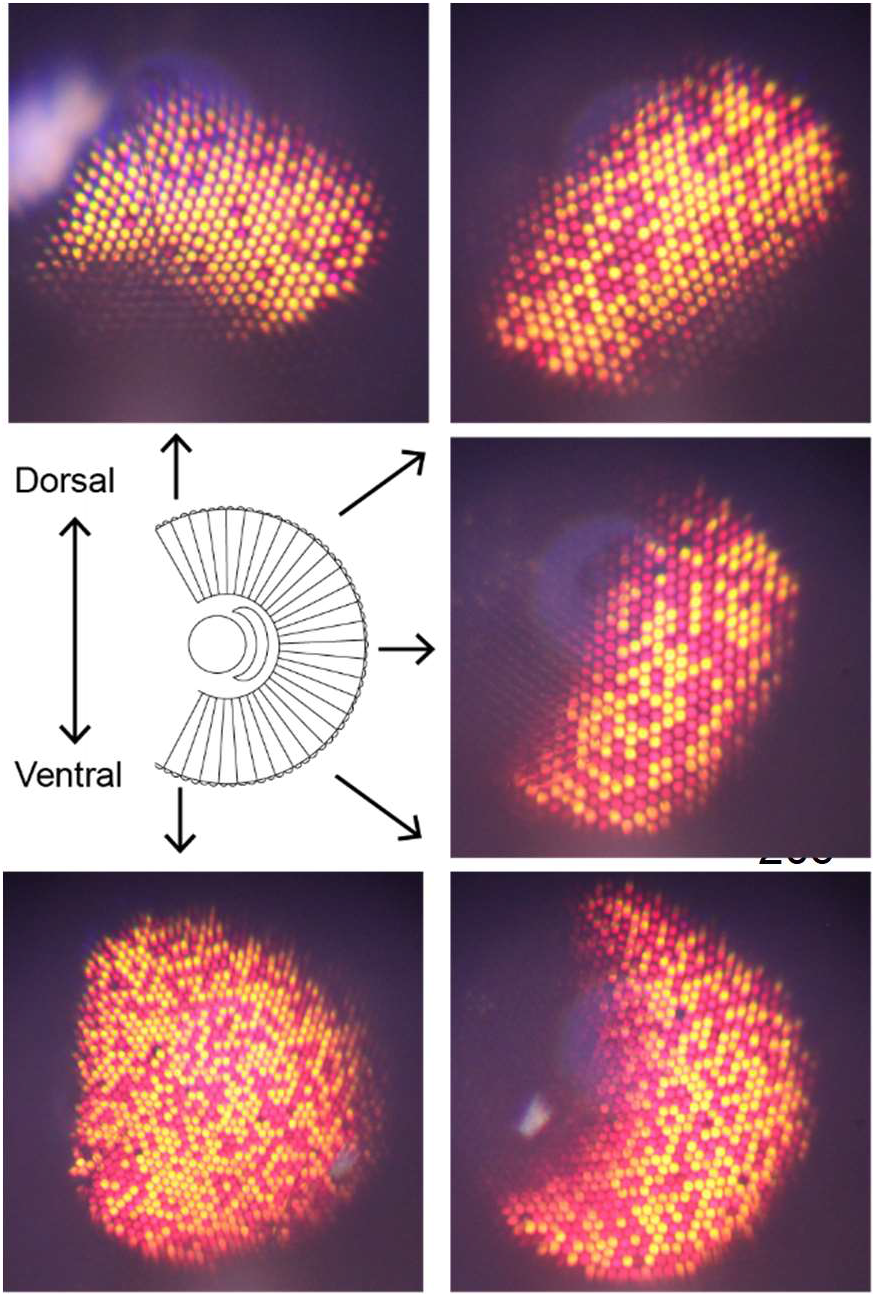
Example eyeshine images. Eyeshine shows which screening pigment is expressed in each ommatidium. We imaged eyeshine across the full dorsal-ventral axis of each butterfly eye. Shown here are a subset of the eyeshine images for an *H. pachinus* male.

In agreement with eyeshine in other *Heliconius* species, every butterfly had red eyeshine indicative of screening pigment expression and yellow eyeshine indicative of no pigment expression (Fig. 4) (Belušič et al., 2021; McCulloch et al., 2016; Stavenga, 2002; Zaccardi et al., 2006). We first measured ommatidia reflectance spectra using a monochromatic camera paired with a series of monochromatic light stimuli (Fig. 5A, see methods). For short wavelengths, yellow ommatidia began reflecting at ~560 nm, which shifted significantly to ~600 nm for red ommatidia (Fig. 5B). Yellow ommatidia also had a peak intensity that was 17.2 ± 13.1% greater than red ommatidia (p < 0.001). The long wavelength cutoff is associated with the reflective tapetum rather than pigment expression (Ribi, 1979) and did not differ between red and yellow ommatidia, with reflectance absent above ~730 nm (Fig. 5B). Reflectance spectra did not vary across groups, but it did vary across the eye (Fig. 5C). Moving across the dorsal-ventral axis of the eye, yellow ommatidia progressively shifted towards longer wavelengths without affecting the shape of the reflectance spectrum (p < 0.05 with Holm-Bonferroni correction). A similar but non-significant shift was observed for red ommatidia (Fig. 5C). Reflectance intensity, in contrast, did not vary with eye region (p = 0.2235).

**Figure 5:**
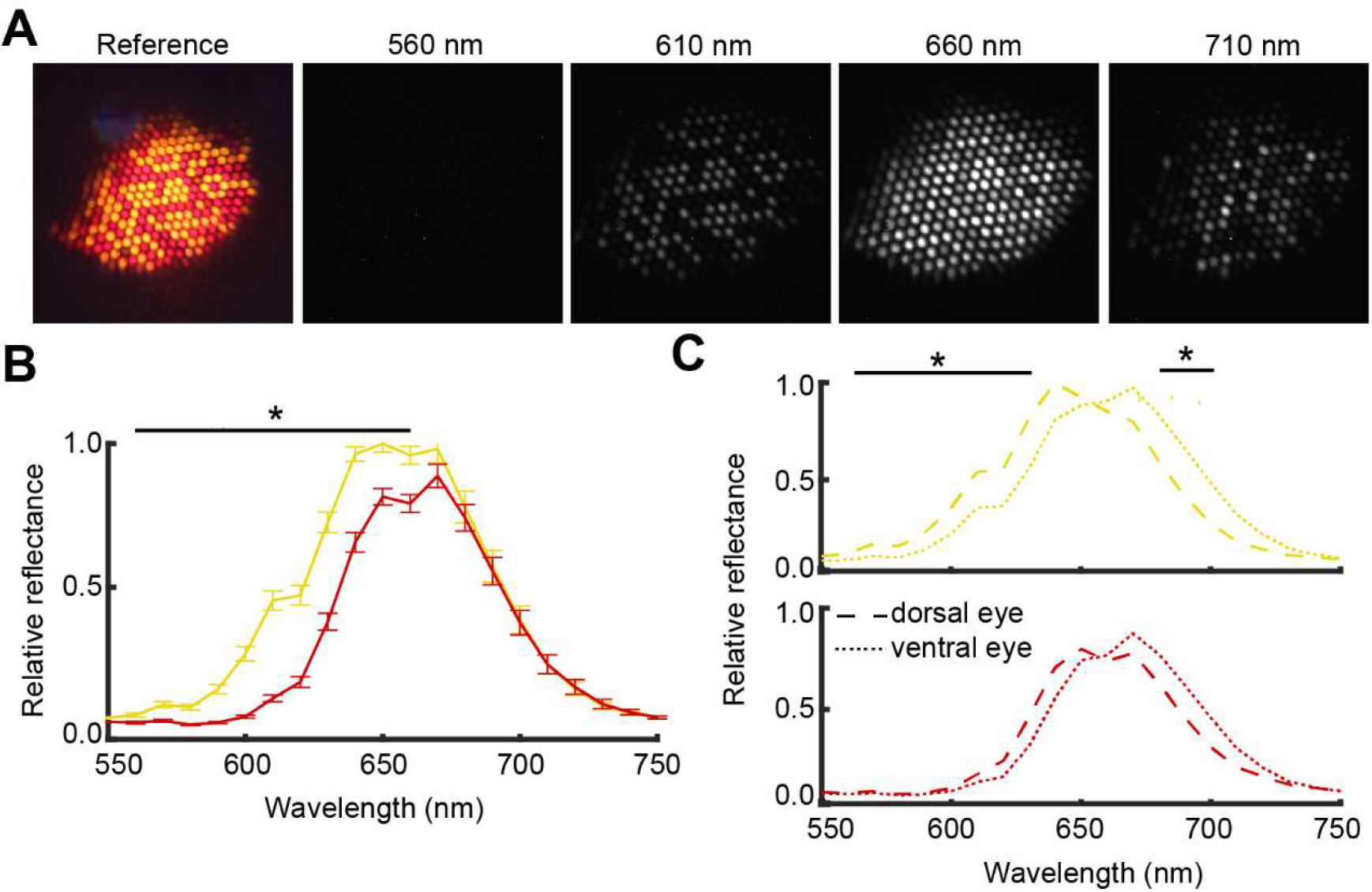
Screening pigment reflectance spectra. (A) The reflectance spectrum of individual ommatidia was measured using monochromatic light and a monochromatic camera. Images shown are for the middle part of the eye for an *H. galanthus* male. (B) Average reflectance spectrum for red and yellow ommatidia (n = 24, split evenly across species and sex, but no F1 hybrids were included). Asterisk indicates where yellow and red reflectance are significantly different (p < 0.05, t-test with Holm-Bonferroni correction). Error bars show mean ± SEM. (C) Red and yellow ommatidia were separated into the dorsal, middle, and ventral part of the eye. The middle part of the eye was intermediate compared to the dorsal and ventral eye and not shown for clarity. Asterisks indicate where reflectance is significantly different across the three regions of the eye (p < 0.05, ANOVA with Holm-Bonferroni correction).

Mirroring the antibody staining results, we also observed differences in the eyeshine distribution between the dorsal and ventral eye. The dorsal eye predominantly had a yellow eyeshine (Fig. 6A), and this did not vary with taxon (F_4,101_ = 1.16, p = 0.33) or sex (F_1,101_ = 3.62, p = 0.06). In contrast, the proportion of yellow ommatidia in the ventral half of the eye (Fig. 6B) varied significantly with taxon (F_4,102_ = 41.76, p < 0.001), sex (F_1,102_ = 127.01, p < 0.001), and the taxon X sex interaction (F_4,102_ = 37.8, p < 0.001). A relatively sharp transition separated the eye into a dorsal ~25% and ventral ~75%, with the decrease in yellow ommatidia occurring across approximately 40 rows of ommatidia.

**Figure 6:**
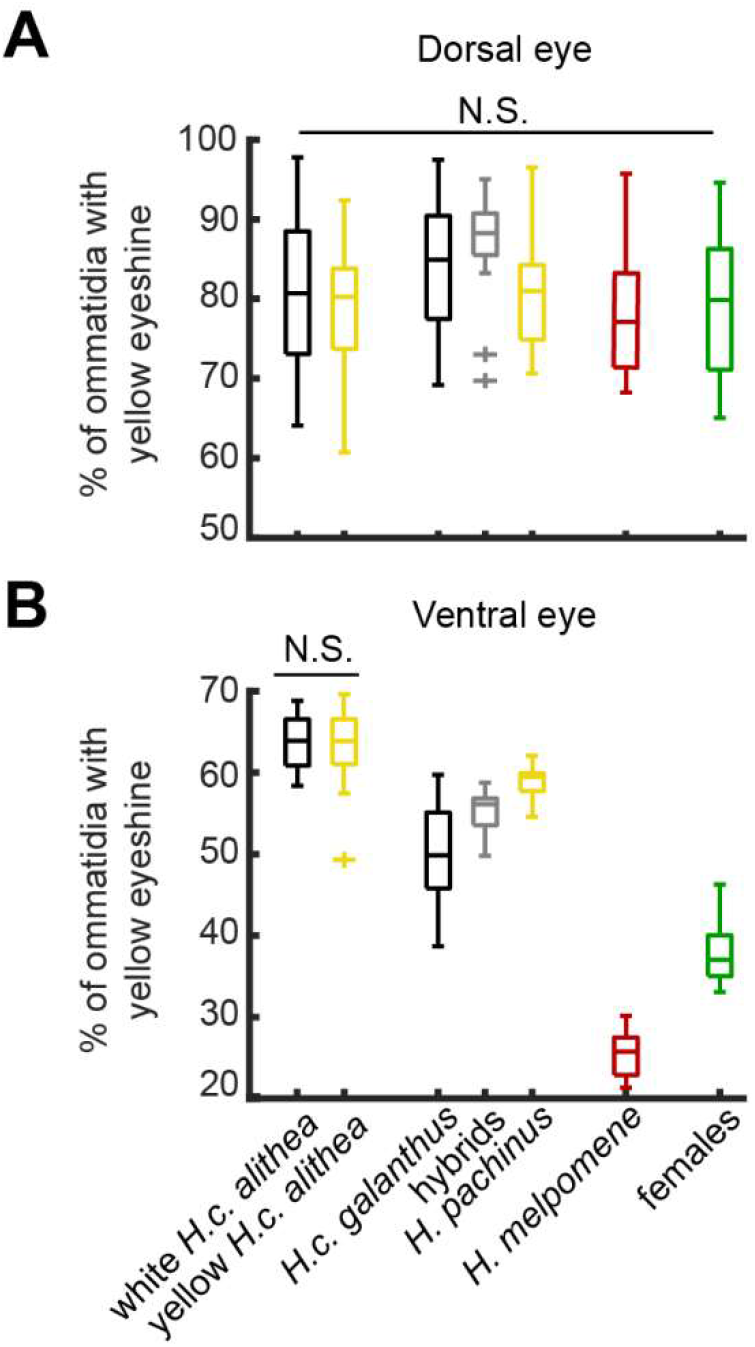
Distribution of eyeshine colors. Boxplots show the proportion of ommatidia that have a yellow eyeshine in the (A) dorsal and (B) ventral eye. Boxes with a taxon name are all males, and all females are grouped together in the final box. No significant differences were detected in the dorsal eye, while all pairwise comparisons except white vs. yellow *H.c. alithea* were significantly different in the ventral eye (n = 15, 15, 15, 14, 14, 15, 15, p < 0.05, t-test with Holm-Bonferroni correction).

For the ventral eye eyeshine, we again observed a sexual dimorphism where male eyes varied and female eyes did not (Fig. 6B). First, female eyeshine was significantly different from males for all taxa (p < 0.001 for all pairwise t-tests with Holm-Bonferroni correction). Further, females did not vary with taxon (F_5,14_ = 0.41, p = 0.83), with 37.9 ± 3.6% of ventral ommatidia having a yellow eyeshine (Fig. 4B). For males, all groups were significantly different from each other (p < 0.05 with Holm-Bonferroni correction) except for *H.c. alithea*, which did not differ between wing colors (yellow eyeshine proportion = 63.5 ± 4.2%, p = 0.90, Fig. 4B). *H. pachinus* (yellow = 58.9 ± 1.9%) had significantly more yellow ommatidia than its sister species *H.c. galanthus* (yellow = 50.5 ± 6.1%), and the hybrid offspring of these two were intermediate (yellow = 55.2 ± 2.5%). In contrast to these *cydno* clade butterflies, *H. melpomene* males had mostly red ommatidia in the ventral eye (yellow = 25.5 ± 2.9%).

Finally, each ommatidial type is typically associated with the same eyeshine color across the eye such that one eyeshine color corresponds to two ommatidial types. However, comparing our antibody staining (Fig. 3B) to eyeshine (Fig. 5B) suggested this relationship differs across taxa (Fig. 7). For *H. melpomene* and females, the proportion of UV-UV ommatidia matched the proportion of yellow eyeshine (Fig. 7A), consistent with results from the dorsal eye. However, this relationship could not explain *cydno* clade males, which had low numbers of UV-UV ommatidia and high proportions of yellow eyeshine (Fig. 7A). Instead, iterating across all possible arrangements, the most parsimonious explanation for these data is that, for all butterflies, UV-UV has a yellow eyeshine and B-B has a red eyeshine (Fig. 7B). In contrast, UV-B differs, with females retaining an ancestral red eyeshine and *cydno* clade males switching to a yellow eyeshine.

**Figure 7.**
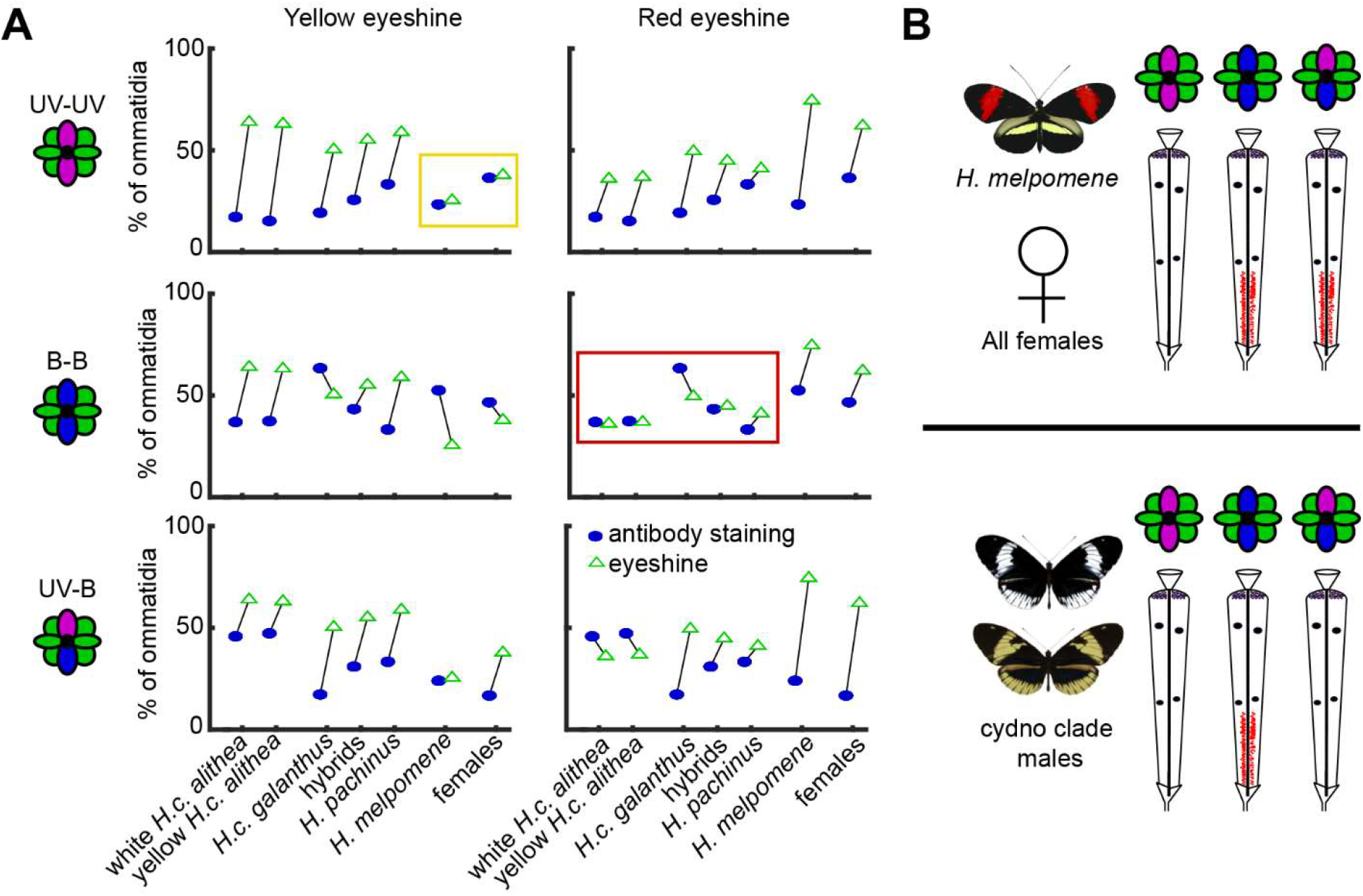
Relationship between screening pigment and ommatidial type. (A) Comparison of eyeshine proportions (columns, Fig. 5b) with each ommatidial type (rows, Fig. 3b). Each panel shows a side-by-side comparison of the proportion of ommatidia with a particular eyeshine color and opsin expression profile. Similar proportions within a panel suggest a one-to-one correspondence across the eye. The two boxes highlight which arrangements parsimoniously minimize the differences for each group. (B) Diagram shows the likely relationship between ommatidial type and eyeshine color. Ancestrally, UV-UV ommatidia have a yellow eyeshine, while B-B and UV-B both have a red eyeshine. *Cydno* clade females likely retain this arrangement, but males appeared to convert UV-B ommatidia from red to yellow.

## Discussion

Overall, our results showed sex-limited variability in eye organization for every metric we examined, with male eyes varying and female eyes appearing similar across taxa. Every butterfly had three ommatidial types (UV-UV, B-B, and UV-B) that matched the inferred ancestral state of all butterflies (Briscoe, 2008), in contrast to the six retinal mosaics detected across all of *Heliconius* (McCulloch et al., 2017). This similarity in the overarching organization of *cydno* clade eyes may be due to a lack of selective pressure to change, but considering the differences we observed, may also suggest this aspect of eye organization is less amenable to rapid evolution. The differences we observed included which UV opsin was expressed, the relative distribution of the three ommatidial types, the distribution of a red screening pigment, and the relationship between ommatidial type and screening pigment.

The first difference in eye organization we detected across these closely related butterflies was which UV opsin was expressed in UV photoreceptors. UV1 is ancestral, while UV2 is a genus-specific adaptation hypothesized to improve the discriminability of a genus specific yellow pigment used for wing coloration (3-hydroxy-dl-kynurenine, 3-OHK) from the yellow pigments used by sympatric, non-*Heliconius* mimics (Briscoe et al., 2010; Bybee et al., 2012). *H.c. galanthus* males strongly prefer to approach and court white females and this was the only taxon where males primarily expressed UV1 (Kronforst et al., 2006). The other *cydno* clade males studied here either prefer yellow females or court both colors equally (Chamberlain et al., 2009; Kronforst et al., 2006), so the observed UV2 expression would serve to enhance conspecific detection in these butterflies.

Many *Heliconius* species have both UV1 and UV2 expressing photoreceptors, but the co-expression of both within single photoreceptors was previously only detected in female *H. doris* butterflies (McCulloch et al., 2017). In contrast to a previous report showing UV1 expression in *H. melpomene* (McCulloch et al., 2017), we detected co-expression of both UV1 and UV2. The most likely explanation for this discrepancy is the antibodies designed for our study had a higher sensitivity, as both our own qPCR (Fig. 2B) and RNA-Sequencing data from the previous report (McCulloch et al., 2017) observed UV2 mRNA at levels ~3X lower than UV1. Since we did not detect UV2 in any females, it is unlikely our results were due to non-specific staining. Co-expression was consistently detected in hybrids that occur rarely in nature, but we also detected this co-expression in a more limited subset of *H.c. alithea* and *H.c. galanthus* males. Other *Heliconius* species have both UV1 and UV2 photoreceptors, and this co-expression may serve a similar adaptive function within the phylogenetic constraint that *cydno* clade butterflies have only three ommatidial types (Finkbeiner and Briscoe, 2021; McCulloch et al., 2017).

Across all of our experiments, we detected a sexual dimorphism where male eyes varied and female eyes did not, suggesting a role in a dimorphic behavior. One possibility is that female eyes are optimized for host plant detection and oviposition behavior. Color vision is important to oviposition in other butterflies, and red sensitive photoreceptors can shift preference towards leaves that appear green rather than yellow to humans (Kelber, 1999; Prokopy and Owens, 1983). Further, all *Heliconius* butterflies specialize on vines in the genus *Passiflora*, suggesting all of the females studied here make similar egg-laying decisions. Nonetheless, female eyes do vary across the genus (McCulloch et al., 2017), which may reflect differences in the specific *Passiflora* species each *Heliconius* species preferentially uses or other selective pressures.

Rather than a role in a dimorphic behavior such as courtship, the differences we observed in males may be related to differences in their natural light environments (Lythgoe, 1979; Sondhi et al., 2021). Males have variable courtship preferences for females with white or yellow wings, but the differences we report here cannot explain this variability (VanKuren et al., unpublished). In particular, white and yellow *H.c. alithea* have different courtship preferences (Chamberlain et al., 2009) but were nearly identical in every analysis. Instead, differences in the relative abundance of different photoreceptor types (Anderson et al., 2017; Bloch, 2015; Fuller et al., 2003), photoreceptor spectral sensitivities (Cummings, 2007; Terai et al., 2006; Torres-Dowdall et al., 2017), and filters functionally similar to butterfly screening pigments (Cronin et al., 2001) have all been linked to differences in light environment in both vertebrates and invertebrates. Here, the ommatidial type distributions (Fig. 3) showed that *H.c. alithea* had the most divergent organization, even compared to the outgroup *H. melpomene*. Additionally, *H.c. alithea* lives in Ecuador, in contrast to all other butterflies in this study which were from Costa Rica, so the observed variability may reflect differences in habitat structure or ambient light levels in these two locations, such as differences in elevation due to the Andes (Dell’Aglio et al., 2022; Sondhi et al., 2021).

Finally, our results suggest an evolutionary change in the relationship between ommatidial type and screening pigment in *cydno* clade males. A direct test of this hypothesis was not possible because the antibody staining protocol washes away the screening pigments. The adaptive value of this change is unclear, but the primary effect should be to decrease the number of red sensitive photoreceptors in *cydno* clade males. This may again be related to adapting *H. cydno* males to the local environment (Cronin et al., 2001; Lythgoe, 1979), while females and red-winged *H. melpomene* males might benefit from the increased number of red sensitive photoreceptors for egg-laying and conspecific detection, respectively.

Comparative studies of sensory systems often show that the periphery can evolve rapidly (Bendesky and Bargmann, 2011), either to support specific behaviors (Auer et al., 2020; Keller et al., 2007) or as an adaptation to the statistics of its natural environment (Fasick and Robinson, 2000; Osorio and Vorobyev, 2008; Regan et al., 2001; Torres-Dowdall et al., 2017; Touhara and Vosshall, 2009). Our results showing several diverse and sexually dimorphic features of eye organization suggest *Heliconius cydno* clade eyes evolved to support both of these adaptive functions. Although less pronounced than the differences observed across distantly related species, our complementary approach of focusing on a group of closely related taxa highlights the value and importance of a zoomed-in view for better understanding visual ecology and the evolution of visual systems.

## Methods

### Animals

The butterflies used in this study were housed in a greenhouse at the University of Chicago that was regularly supplemented with new butterflies from breeders located in Ecuador (H.c. *alithea*) and Costa Rica (H.c. *galanthus* and *H. melpomene). H. pachinus* and hybrids were reared in Panama and transported to the University of Chicago for experiments. All butterflies were at least 3 days old at the time of experiments.

### Antibody staining

Butterflies were decapitated into 0.01 M phosphate buffered saline (PBS) where eyes were dissected using forceps. Eyes were fixed at room temperature for 15 minutes in 4% paraformaldehyde in PBS. Fixed eyes were cryoprotected in a 25% sucrose in PBS solution overnight at 4°C. Eyes were then frozen in Tissue Tek O.C.T., sectioned at 14 μm on a cryostat, and placed on slides to dry overnight.

Cross sections of the distal eye were immunostained with antibodies specific to blue and UV sensitive opsins. The anti-blue opsin antibody was generated against the peptide INHPRYRAELQKRLPC in rabbits and was a gift from Michael Perry (Perry et al., 2016). Since *Heliconius* butterflies have both a UV1 and UV2 opsin (Briscoe et al., 2010), we generated new antibodies specific to each (GenScript). For UV1, the antibody was generated in guinea pigs against the peptide GLDSADLAVVPEC. For UV2, the antibody was generated in mouse against the peptide GLSSAELEFIPEC. To stain sections, slides were first washed in chilled acetone for 5 minutes, 2X10 minutes in 0.01 PBS, 2X10 minutes is 0.3% Triton X-100 in 0.01 M PBS (PBST), 1X5 minutes in 1% sodium dodecyl sulfate in PBST, and 3X10 minutes in PBST. Slides were then blocked for 1 hour in 1% bovine serum albumin in PBST. Primary antibody was applied overnight at 4°C in 1:300 dilutions. The following day, slides were washed 5X10 minutes in PBST before applying the secondary antibody. Secondary antibodies (Abcam) were diluted 1:2000 in blocking solution and applied to the slides for 2 hours at room temperature. These antibodies were goat anti-rabbit Alexafluor 488, donkey anti-guinea pig 555, and donkey anti-mouse Alexafluor 647. After staining, slides were finally washed 5X10 minutes in PBST and stored in Polymount (Fisher Scientific). Eye slices were imaged using a Zeiss LSM 510 confocal microscope using a 20X objective.

### Quantification of ommatidial types

We quantified the distribution of the three ommatidial types by counting the number of UV-UV, B-B, and UV-B ommatidia in each slice with an automated program. We first generated three binary masks for each slice, with one for UV staining, one for blue staining, and one for the merged image. Ommatidia were automatically identified using the MATLAB function *bwareafilt* on the binary merged image. We overlayed the ommatidium boundaries on the binary UV and blue images and defined the ommatidium as UV or blue positive if at least 15% of the pixels were stained for the opsin. This threshold minimized variability, but results were not qualitatively affected by different values.

We controlled for the quality of our automated program in two ways. First, we visually counted ommatidia in 12 sections and compared results, finding less than 4.9% differences in ommatidial type across all sections. Second, we averaged the measurements across 2-4 sections per eye. Across all eyes, the proportions of each ommatidial type differed by an average of 3.8 ± 3.7%.

To compare the distributions across groups, we used hierarchical clustering of the ommatidial types. Each of the three types were used as different dimensions, with each individual as a unique data point. We clustered based on the Euclidean distance using an average linkage function, which maximized the cophenetic correlation (r = 0.77). Results and conclusions were not affected when using alternative distances or linkage functions.

### qPCR

Eyes were dissected from a butterfly and immediately placed in RNA-later and stored at −80°C. Prior to RNA extraction eyes were repeatedly washed in PBS. RNA was extracted and converted to cDNA using a Qiagen RT-PCR kit. Expression levels for UV1 and UV2 were assayed using SYBR green. UV1 primers were 5’-CGCTCACTGTGTGCTTCCTCTT-3’ and 5’-AGTCTTGCAAGCTACCGCGG-3’. UV2 primers were 5’-TACCGTGTGCTTCCTTTATGTTG-3’ and 5’-ACCCTTGCAAGCGATCGCAG-3’.

### Eyeshine

Eyeshine images were collected using a custom built epi-fluorescent microscope following a published design (Stavenga, 2002). White light (DH-2000S, Ocean Optics) entered the microscope vertically where the beam was expanded to fill the imaging objective using two lenses placed confocally (40 and 80 mm, Edmund Optics). A half-silver mirror directed the light through a 20X, 0.4 NA objective (Zeiss LD-Plan-Neofluar) that was focused on the eye of a butterfly. After reflecting off the tapetum at the base of an ommatidium, light re-entered the horizontal arm of the microscope where it was magnified using 80 and 20 mm lenses placed confocally with each other. The eyeshine was then photographed using a digital camera equipped with an infinity focused lens (Canon EOS Rebel T5).

For each experiment, a butterfly was restrained in a custom collar with beeswax and placed on a rotating platform near the focal point of the imaging lens. The eyeshine was brought into focus using three linear actuators. The butterfly was dark adapted for at least one minute before each image. After each image, the butterfly was rotated to a new, non-overlapping position along the dorsal-ventral axis of the eye.

We quantified the eyeshine distribution by counting the number of red and yellow ommatidia in each photo. Each image was analyzed blind to taxon, sex, and location along the dorsal-ventral axis of the eye. A randomly selected 20% of the eyeshine images were included twice to ensure repeatability of the count, finding a maximum of a 2.6% difference in the proportion of yellow ommatidia counted. To calculate the proportion of yellow ommatidia in the dorsal eye, we combined the two dorsal-most images. For the ventral eye, we combined counts from the ventral half of the photos, rounded down. Results were not affected combining different numbers of photos.

### Eyeshine spectral reflectance

We measured the spectral reflectance of individual ommatidia using the same epi-fluorescent microscope. After taking a reference image, we then rerouted the white light through a monochromator (MonoScan-2000, Ocean Optics) and replaced the digital camera with a monochromatic camera (Prosilica GX1050, Allied Vision Technologies). Measuring an accurate reflectance spectrum required controlling light intensity across different wavelengths. We first used neutral density filters (Thorlabs) to minimize differences in the number of photons per second that entered the microscope. We then scaled the shutter time for each wavelength such that each exposure would contain the same number of photons. Tests using a mirror in place of a butterfly showed this procedure was effective at equalizing photon flux.

The reflectance spectrum was measured from 550 to 750 nm in 10 nm steps. Preliminary experiments showed no reflectance outside this range. After orienting the butterfly, we bleached the eye with white light for 10 minutes. Each stimulus was 1.0 × 10^15^ photons, which was an average shutter time of 6.6 ± 1.1 seconds. We performed control experiments where we compared images collected at the beginning and end of a 10-minute exposure, which confirmed that the low intensity of monochromatic light was unable to induce corneal adaptation. For each butterfly, we measured the spectral reflectance for an image from the dorsal, middle, and ventral part of the eye.

We analyzed the spectral reflectance of individual ommatidia using ImageJ (Schindelin et al., 2012). Images were imported as a z-stack, which allowed us to manually select each ommatidium as the same region of interest across photos. Each ommatidium was identified as red or yellow by overlaying the reference eyeshine image. The reflectance at each wavelength was then defined as the average pixel intensity within the selected region of interest. Images were 8 bit (0-255), and ommatidia were excluded from further analysis if the peak intensity was less than 125 or greater than 250.

## Acknowledgements

We would like to thank Daniel Baleckaitis for his help with immunostaining, Laura Southcott for collecting and breeding animals, and Doekele Stavenga and Primoz Pirih for valuable discussions and technical development.

## Data Accessibility

All of the data presented here are available from the Dryad Digital Repository at doi: https://datadryad.org/stash/dataset/10.5061/dryad.5hqbzkh7v.

